# A Role for Gαz in regulating seizure susceptibility

**DOI:** 10.1101/567628

**Authors:** Rainbo Hultman, Okechi Boms, Stephen Mague, Dalton Hughes, Victor Nadler, Kafui Dzirasa, Patrick J. Casey

**Affiliations:** Departments of Biochemistry Pharmacology and Cancer Biology Duke University Medical Center, Durham, North Carolina 27710; Psychiatry and Behavioral Sciences and of Pharmacology and Cancer Biology Duke University Medical Center, Durham, North Carolina 27710; Pharmacology and Cancer Biology Duke University Medical Center, Durham, North Carolina 27710

**Keywords:** Gnaz, seizure susceptibility, G protein, epilepsy

## Abstract

Much about the molecular mechanisms underlying seizure susceptibility remains unknown. A number of studies have indicated that the neurotrophic factor BDNF plays an important role in mediating seizure susceptibility. Recently, we found that the heterotrimeric G – protein, Gz, which is known to endogenously couple to monoaminergic receptors, such as serotonin, norepinephrine and dopamine receptors, regulates BDNF-induced signaling and development in cortical neurons. Interestingly, several of the receptors that Gz endogenously couples to have also been shown to be associated with seizure phenotypes (5HT_1A_-serotonin and D2 dopamine). Here we characterized seizure susceptibility in Gz-null mice, behaviorally and electrographically, finding that Gz-null mice have increased seizure susceptibility using a modified version of the pilocarpine model of status epilepticus. Local field potential (LFP) data recorded from six brain regions-amygdala, dorsal hippocampus, ventral hippocampus, motor cortex, somatosensory cortex, and thalamus-showed robust electrographic seizure activity for Gz-null mice compared with low or no seizure activity in wild-type controls.

## Introduction

Seizures are the behavioral and electrophysiological manifestation of epilepsy ^1^, and are also found in sudden unexpected death in epilepsy (SUDEP) ^2^ and autism ^3^. Epilepsy causes adverse psychiatric, physical and social harm; alone, it accounts for 0.5% of the global burden of disease ^4^. Much about the mechanisms underlying seizure disorders are still not known; current therapies have some success such that about 65-70% of temporal lobe epilepsy cases can be normalized, at least initially, but there is a great need to identify causes and treatments that address the underlying brain dysfunction in order to improve long-term efficacy.^5,6^

A better understanding of the molecular pathways underlying seizure disorders is essential for the development of better treatments. Several pieces of evidence suggest that neurotrophins, and BDNF in particular, contribute to seizure susceptibility. BDNF has been shown to be upregulated after limbic seizures, particularly in the dentate granule cells of the hippocampus in mouse models ^7–10^, and an increase in BDNF protein levels has been identified in hippocampal specimens from patients with temporal lobe epilepsy ^11^. Preclinical models manipulating BDNF activity have altered development of epilepsy ^12^ ^13^ ^14,15^. Taken together, these studies indicate that pathways implicated in BDNF regulation are crucial for mediating seizure susceptibility.

Previously, we demonstrated that the heterotrimeric G protein, Gz, negatively regulates BDNF-stimulated signaling with regard to axon development in cortical neurons ^16^. Gz is a member of the Gi subfamily of heterotrimeric G proteins, and like others of this class, is inhibitory toward adenylyl cyclase. It has also been shown to impact downstream effectors Rap1GAP and Eya2^17,18^, and its activity can be regulated by the regulator of G protein signaling protein RGS20^19^. There are several interesting biochemical properties that distinguish Gz from other Gi subfamily members; Gz has an extremely slow kinetics with regard to GDP-dissociation and GTP hydrolysis (~200-fold slower than other G□i subunits)^20^, and is the only member of the Gi subfamily to be insensitive to C-terminal ADP ribosylation by Pertussis toxin ^20,21^. Another trait that distinguishes Gz from other Gi subfamily members is that Gz expression is restricted to vertebrates and is predominantly localized to the CNS ^22^. Gz plays a crucial role in neuronal signaling, particularly with regard to monoamine neurotransmitter signaling ^22,23^.

In contrast to findings in Gi-null mice, which have more severe impairments across multiple physiological systems, Gz-null mice have demonstrated almost exclusively neuronal and neuroendocrine phenotypes, and largely only in response to pharmacological interrogation ^24^. Such studies demonstrate that in vivo, Gz endogenously couples to serotonin 5-HT_1A_^25,26^, dopamine D2^27^, □_2A_-adrenergic^28,29^, □-opioid^30^ and EP3 E-prostanoid receptors^31^. Accordingly, Gz-null mice have demonstrated an increased anxiety behavioral phenotype^25,26^; Gz-null mice have a failure to respond to serotonin receptor 5-HT_1A_ agonist 8-OH-DPAT, with regard to both anxiety and CA1 pyramidal neuron K^+^ current^26^. Gz-null mice are also insensitive to the locomotor inhibition effects and alterations in the levels of pituitary adrenoccorticotropin releasing hormone (ACTH) induced by the D2 agonist, quinpirole^27,32^. Additionally, Gz-null mice have altered pre-pulse inhibition (PPI) responses, a process that is highly dependent upon the D2-dopamine receptor^33^. Several of these monoamine neurotransmitter systems have been linked to seizure susceptibility as well ^34–37^. Gz is thus implicated in several pathways, both directly and indirectly, that are linked to seizure susceptibility. Given these findings, we hypothesized that Gz would have a negative regulatory impact on seizure susceptibility, much in the same way that it impacts BDNF-related signaling, and set out to test this hypothesis.

## Results

In order to test the hypothesis that Gz negatively impacts seizure susceptibility, we implemented a well-characterized seizure induction model in conjunction with Gz-null mice. Several animal models have been developed to emulate status epilepticus (SE)^38^, a continuous state of seizure activity. Most animal models of SE (and epileptogenesis) involve either a chemical or electric stimulant that triggers convulsions; however, the manifestation of SE indicates that the brain has been triggered beyond this initial stimulus into a recurring feedback loop of stimulation. Such continuous activity is recordable by EEG and observed behaviorally by the absence of consciousness and the occurrence of motor seizures ^39^; tonic, clonic seizures may persist for hours ^40^. Notably, this same approach is used to monitor seizure activity in patients with epilepsy^41^. In order to determine whether Gz-null mice had altered seizure susceptibility, a low dose of pilocarpine (pilocarpine), was employed in the first part of the study. Pilocarpine is a muscarinic acetylcholine receptor agonist that is widely used to induce status epilepticus ^39^. The culmination of a number of observations by different groups indicate that pilocarpine activates M1 muscarinic receptors in the brain, causing an initial imbalance between excitatory and inhibitory transmission leading to the onset of status epilepticus; status epilepticus is thus maintained long after the action of pilocarpine by this imbalance ^39^. Another advantage to a pilocarpine model is that it induces development of brain lesions that mimic those found in patients with temporal lobe epilepsy, thus providing some evidence that it mimics some of the underlying causes of epileptogenesis, and not just the symptomatic behavior ^39^.

### Behavior

All behavioral studies detailed herein were conducted blind to genotype. Low dosages of pilocarpine (150-250mg/kg) were used that were not expected to induce status in normal mice. Mice were observed and their behavior was recorded at 5 min intervals for 4 h. Using such behavioral measures, we found that Gz-null mice demonstrated significant seizure susceptibility in response to a subthreshold dose of pilocarpine, where wild-type mice had few low-grade or no seizures. The Gz-null mice repeatedly demonstrated more severe seizure behavior than WT controls (Table 1, Fig 1A). Mice lacking Gz also had a significantly shorter latency to onset of seizures than the WT animals (Fig.1B).

**Table 1.**
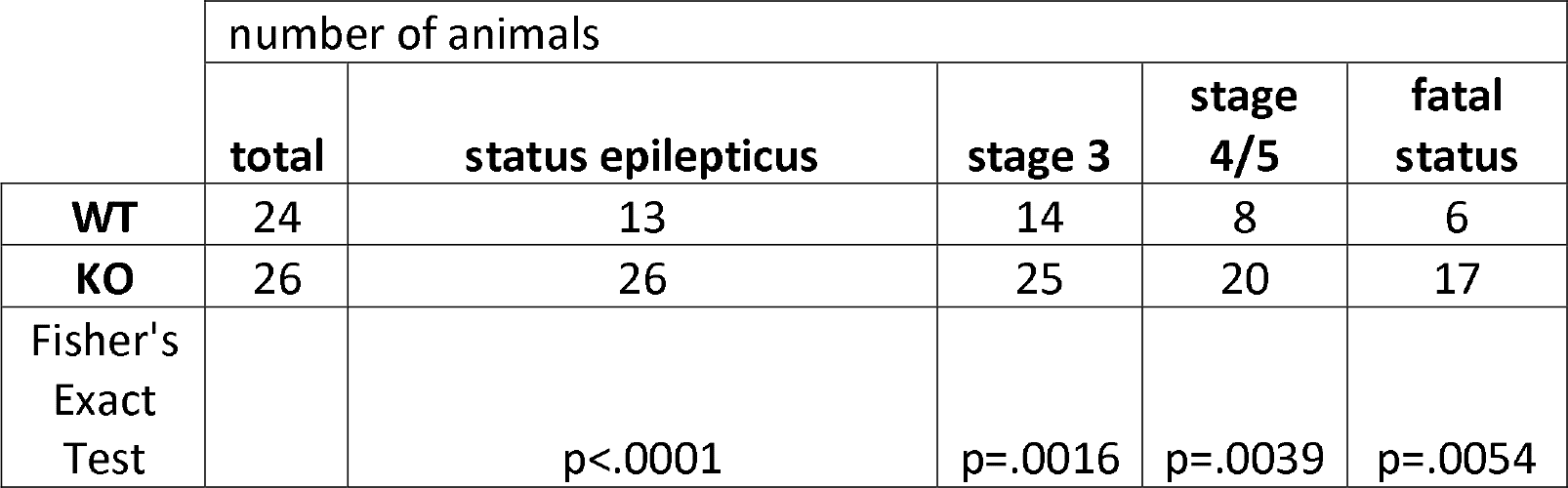
Gz-null mice demonstrate increased seizure severity. Male BALB/c mice, ~16 weeks old were treated with 150-250 mg/kg pilocarpine and monitored for seizures. Status epilepticus (SE) was defined as a continual state (>30 min) of seizure of class 2 or higher, with classifications defined as: stage 0, stopping, staring; stage 1, facial stereotypes such as lip smacking or foaming at the mouth, as well as loss of conscious response to sounds and smells; stage 2, compulsive head nodding; stage 3, forelimb clonus (rapid shuffling of paws); stage 4, seizure including rearing onto hind limbs; stage 5, seizure where mice roll over or throw their bodies around the cage; fatal status (FS) seizure terminating in death. Numbers of mice achieving stages: SE, stage 3 seizures, stage 4 or 5 seizures or fatal status are indicated alongside Fisher’s Exact Test p-values.

**Fig. 1.**
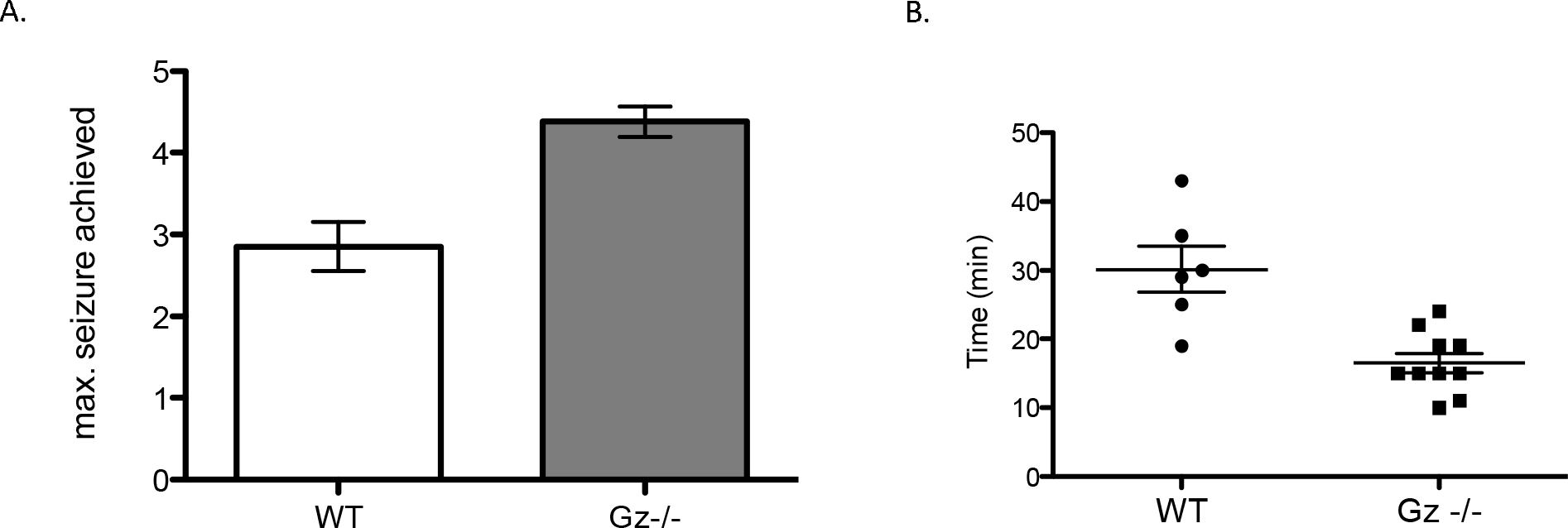
Gz-null mice have decreased latency to seizure onset. A) average maximum class of seizure achieved; statistical significance is shown as p<.001 (***). B) latency to the onset of status epilepticus in response to 250 mg/kg pilocarpine. Latency is plotted only for animals that achieved status. Error bars indicate SEM. Statistical significance is shown as p<.001 (***).

### Electrographic Seizure (EGS) Onset and Spread

We next sought to understand the electrographic characteristics and neural circuitry of seizures in Gz-null mice. To do this we implanted electrodes in six brain regions in wild-type and Gz-null BALB/c mice and measured the activity (local field potentials) in multiple brain regions in-vivo in freely behaving rodents. Wild-type (WT) and Gz−/− mice were implanted with electrodes in regions that have been shown to be of importance to limbic seizures, including dorsal and ventral hippocampus, medial anterior thalamus, amygdala, and motor and somatosensory cortex. After a two week recovery, these animals were treated with a low dose of pilocarpine (180 mg/kg) and neurophysiological EEGs were recorded in all six brain regions prior to seizures, upon induction of seizure and through each stage as each progressed through the various seizure stages (Figure 2). The duration of the experiment was approximately 3-3.5 hours for all four trials, dependent on the duration of seizure. Following the five-stage behavior scale already described, all four Gz−/− mice demonstrated more severe seizure-like behaviors than the 4 WT animals. Two of four Gz−/− mice reached stage 5, with barrel rolling. All four Gz−/− mice eventually died in status (fatal status) during the electrophysiological recordings.

**Fig. 2.**
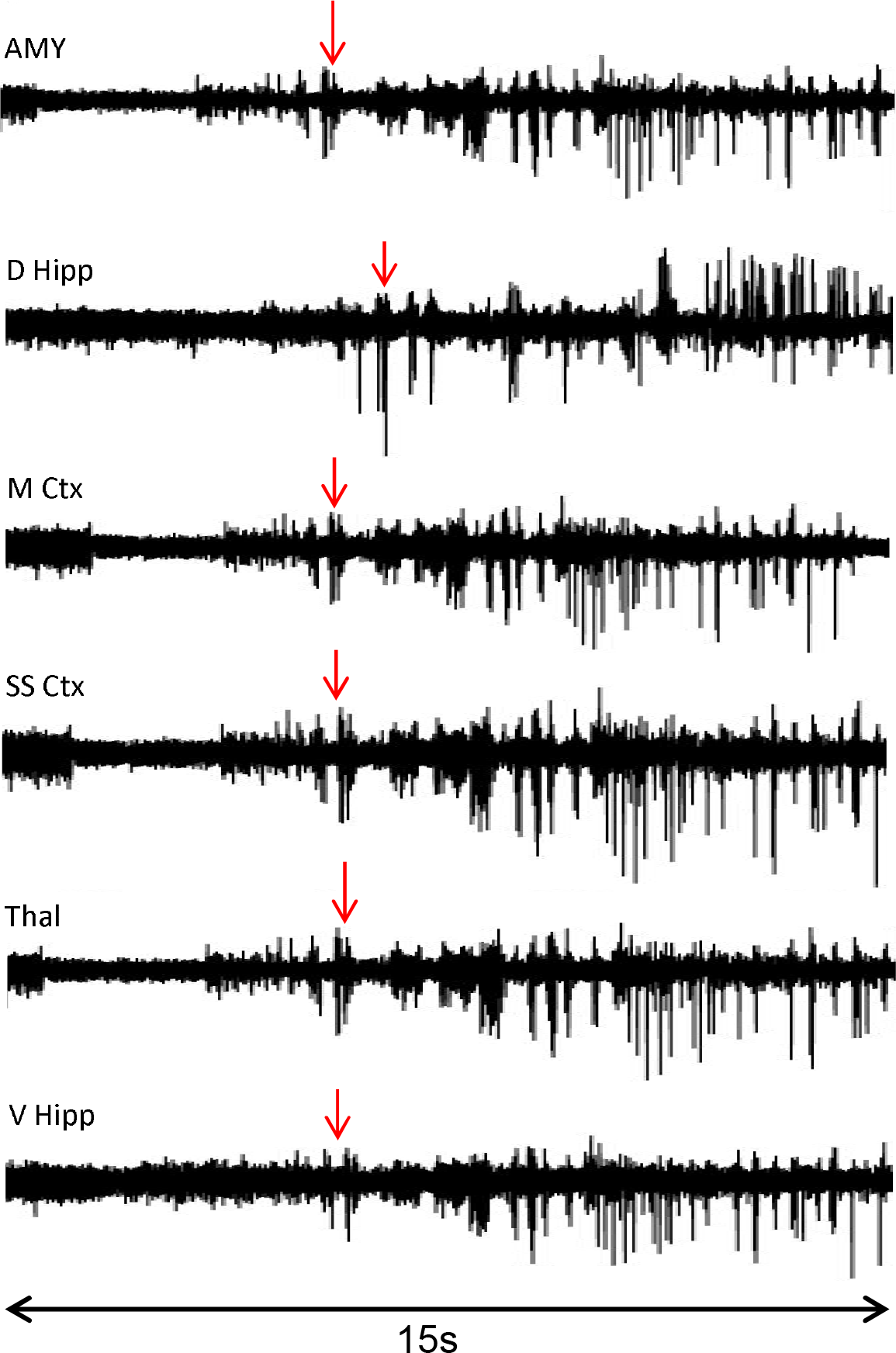
Representative electrographic seizure identification. The onset of seizure is represented by the change in LFP excitability (mV); red arrows show the wave spike identified as the onset of the seizure. LFP data shows a reduction in baseline activity that transition into recurrent, increased spiking for at least 10 seconds. Representative examples provided are from seizures in a Gz-null mouse.

On average, the onset of electrographic seizures was earlier in Gz−/− mice when compared to WT mice, and two WT mice had no electrographic seizure activity in the regions recorded. The average time of first EGS, after pilocarpine injection, for Gz−/− mice was always much lower than that in WT mice (e.g. 4.9+/-1 min. in ventral hippocampus of Gz−/− compared with 28.7+/− 4min. in that of WT animals, Fig. 3). Within the Gz−/− group, there was variation in the seizure spread pattern.

**Fig. 3.**
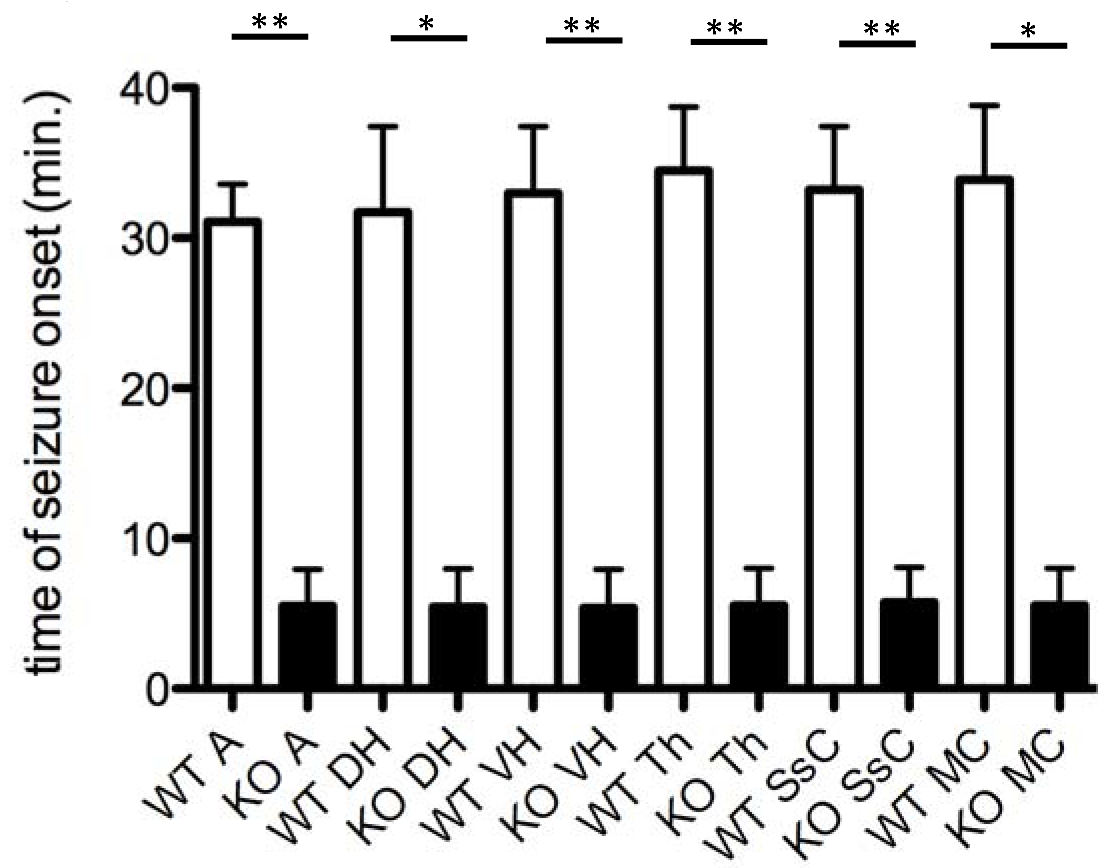
Electrographic seizure onset. The average time of electrographic seizure onset (minutes) for each recorded brain region for the WT and Gz knockout (KO) groups. Brain region abbreviations and p-values: Amygdala (A), ** p=.006, dorsal hippocampus (DH), * p=.0164, ventral hippocampus (VH), ** p=.0097, thalamus (Th), ** p=.0076, somatosensory cortex (SsC), ** p=.0081, motor cortex (MC), * p=.01.

### Characteristics of Gz−/− Seizure

Across all regions, the EGS in Gz−/− mice were more robust and severe than WT mice (Figure 4). In addition, the seizures in Gz−/− mice showed a pattern of progression that was roughly divided in three phases (Figure 5). In Phase 1 (P1), the LFP waves showed seizure patterns that mark the onset of mild seizure activity; the magnitude baseline voltage is similar to that before pilocarpine injection. Phase 2 (P2) was marked by a decrease in LFP baseline, followed by a pattern of increased LFP amplitude. In Phase 3 (P3), there was a significant increase in the magnitude of baseline voltage compared to P1 and P2; two out of four Gz−/− mice died in P3.

**Fig. 4.**
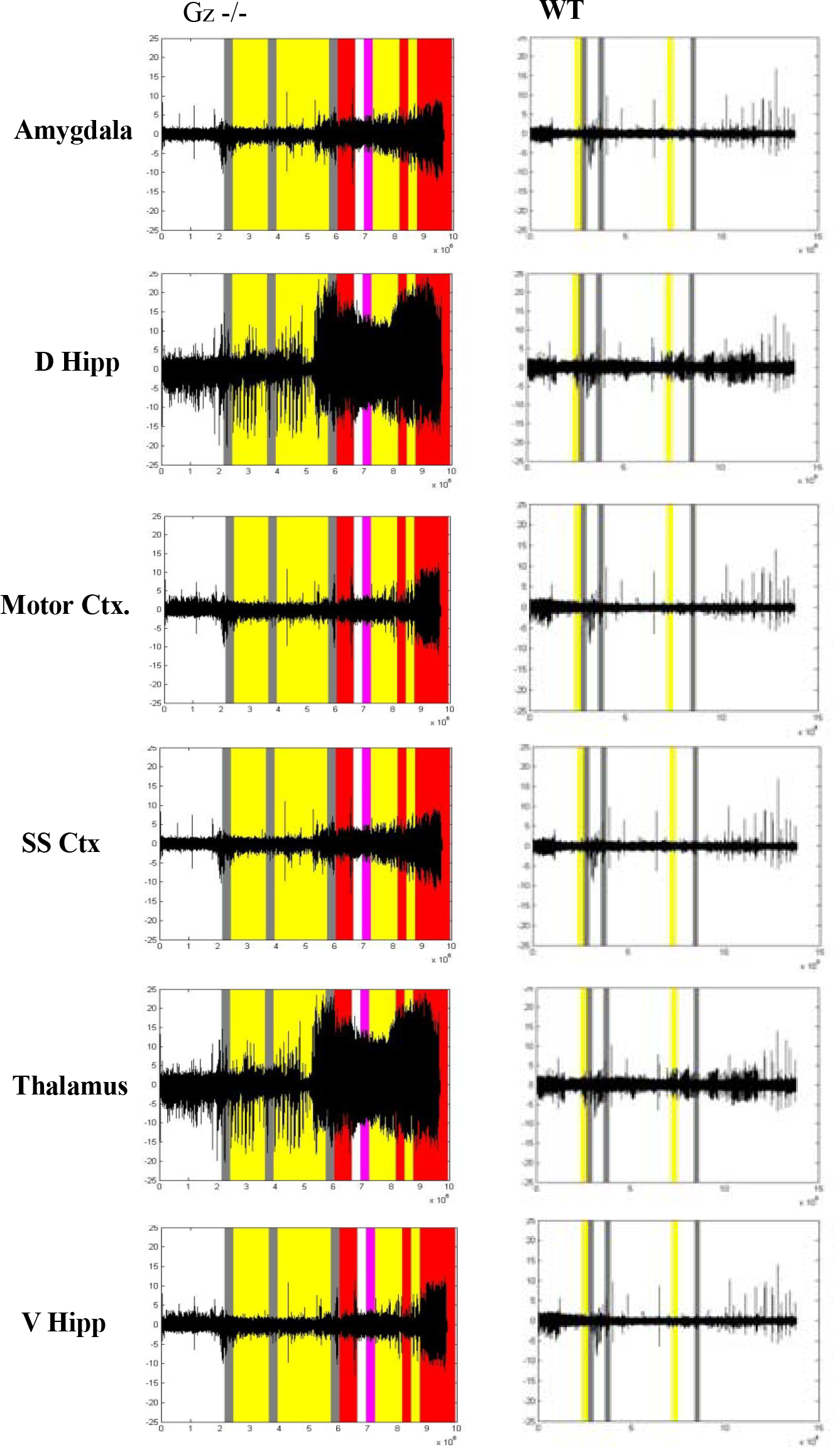
Electrographic seizure activity corresponds to seizure behaviors. Representative examples of littermates of each genotype are shown. Gz−/− mouse (left) showed more robust LFP activity and severe seizure behavior than WT animals (right). The plot shows change in voltage (mV) on the y-axis over the duration of the experiment (time in ms on the x-axis). The behavioral seizure scale (1-5) was color coded to relate the severity of behavior during recordings: 0-white; 1-white; 2-gray; 3-yellow; 4-magenta; 5-red (refer to Methods for a description of corresponding behavior).

**Fig. 5.**
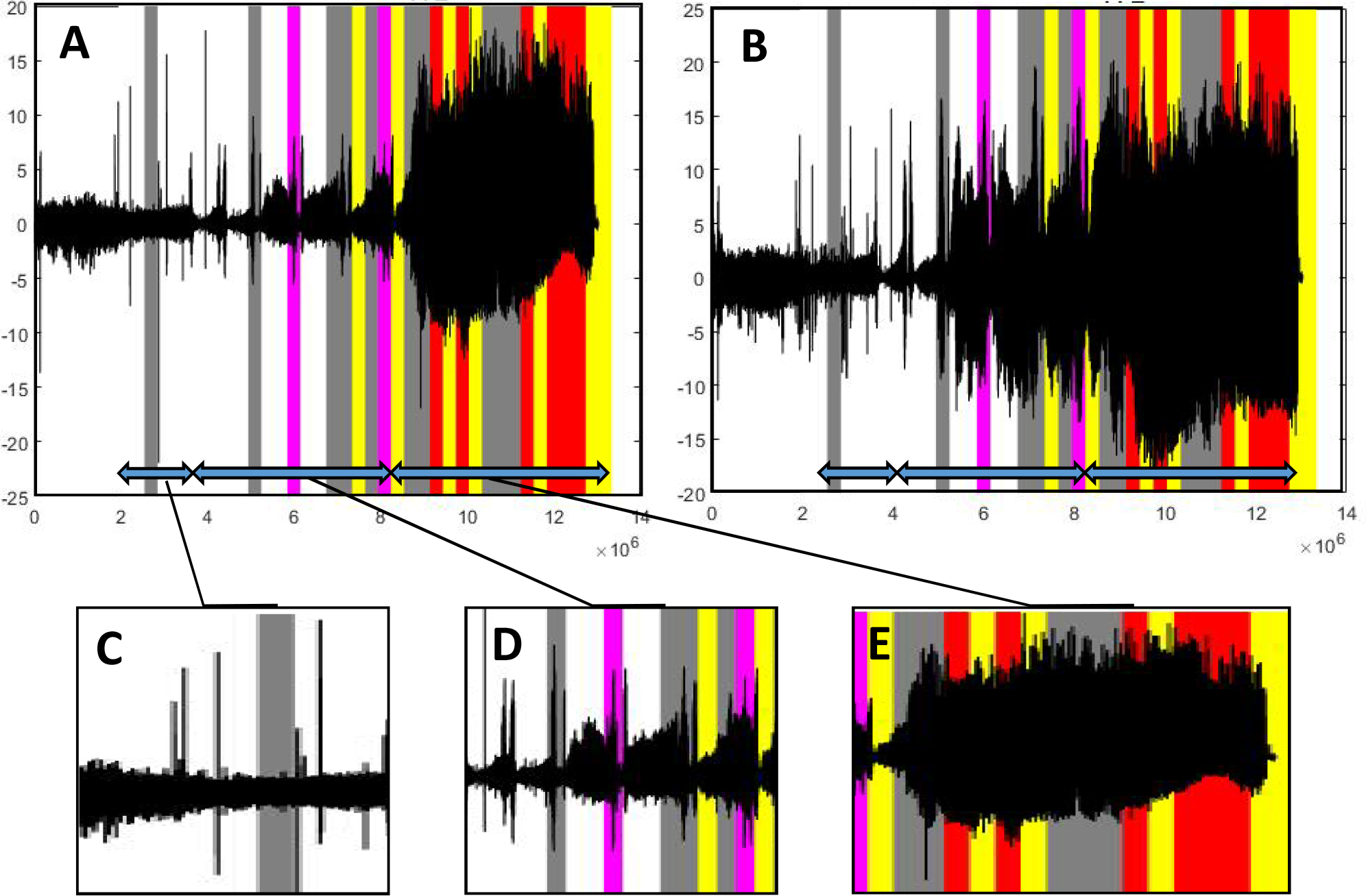
Phases of Gz-null electrographic seizures. Representative examples of the three phases, P1, P2, and P3, of Gz−/− electrographic seizures are marked by double headed arrows in the A.) dorsal hippocampus and B.) ventral hippocampus. The plots show change in voltage (mV) on the y-axis over the duration of the experiment (time in ms on the x-axis). C.) Phase 1 of the Gz−/−EGS D.) Phase 2 was marked by progressive increase in LFP amplitude followed by a sharp decline to baseline levels and E.) Phase 3 showed a rapid and sustained increased in LFP amplitude.

## Discussion

Here we have shown that removal of the Gαz protein increased seizure susceptibility both behaviorally and electrographically in a mouse model. The Gαz-null group demonstrated substantially more severe seizure behavior, which was measured as a higher count of stages 3, 4, and 5 when compared to the WT group. The LFP data support this finding, as evidenced by Gαz-null mice having earlier and more robust changes in LFP activity, corresponding well with behavioral differences (Figure 4). Stage 4 and stage 5 behaviors, only observed in the Gαz-null animals, were more likely to happen during phase 3 when there was a substantial increase in LFP amplitude. This distinction is particularly noteworthy as these advanced stages are considered generalized seizures and may lead to further work in characterizing the mechanisms of hyperexcitability in Gz-null mice. In the WT group, only one of three mice showed signs of electrographic seizures. In WT mice that did not have electrographic seizures, there were irregular patterns after pilocarpine injection that were not considered seizure because they did not last for at least 10 seconds.

These results showed that Gz-null mice had more severe behavioral and electrographic seizures and faster spread of seizures across all of regions that were measured, compared with WT mice. The behavioral and electrographic results confirm the role of Gαz in mediating seizure severity and susceptibility. The site of seizure onset and progression for the WT was the hippocampus, which closely matches the pattern from other studies^42^; the Gz-null mice showed several different patterns, with >10sec seizures of several animals originating in the hippocampus as well and one in the amygdala.

Interestingly, the GNAZ gene is found at chromosomal site 22q11.2, a region with extremely common microdeletions and microduplications associated with severe behavioral abnormalities^43^. One study found that a patient with gene duplications in the GNAZ region has seizures and epilepsy, suggesting that our findings may reflect a naturally occurring biological phenomenon^44^. This same patient also has psychosis and anxiety, which also overlap well with phenotypes previously described in Gz-null mice^44^. It has been shown that seizure disorders have a high comorbidity with other psychiatric disorders, such as personality disorders, depression and anxiety disorders ^45,46^. Interestingly, Gz-null mice have been shown to display heightened anxiety and depression-like behavior ^25^ and show an altered response to psychoactive drugs such as amphetamine ^32^. Further studies into the mechanisms by which Gz-null mice have increased seizure susceptibility, may result in finding a common mechanistic linkage.

There are several cell signaling networks in which Gz plays a key role that have also been demonstrated to contribute significantly to seizure susceptibility. Gz has been linked to signaling by the neurotrophin, BDNF, which has also been shown to play a major role in mediating seizure susceptibility and severity ^6,14,16^. Studies have shown that BNDF is upregulated in areas implicated in epileptogenesis, and interference with BDNF signal transduction inhibits the development of epilepsy ^47^. Mice heterozygous null for BDNF display a delay in epileptogenesis by use of the kindling model of temporal lobe epilepsy ^12^. The findings in this current work further implicates BDNF expression in epileptogenesis.

Gz has also been shown to couple in vivo to the major monoamine neuromodulatory systems, serotonergic, dopaminergic, and adrenergic, which also play important roles in seizure susceptibility ^22,37^. In particular, the 5-HT_1A_ serotonin receptor has been shown to increase seizure threshold^48^. When compared to WT littermates, 5-HT_1A_ knockout mice display increased anxiety-related behaviors across ethological behavioral tests ^49,50^. Similarly, the D2-dopamine receptors play an important role in mediating anticonvulsant effects within the context of acute pilocarpine-induced seizure models^51^. While other G□i subunits have been shown to couple to metabotropic glutamate receptors, chimeric studies indicate that G□z binding to mGluRs is much more limited ^52,53^; Gz has not been found to couple endogenously to mGluRs ^24,52,53^. Given the complex relationship between serotonin and dopamine in epilepsy, a role for Gz in mediating seizure susceptibility raises the possibility of Gz-regulated cell signaling events downstream of these neuromodulators in regulating seizure susceptibility.

## Materials and Methods

### Animal Care and Use

All experiments were carried out in accordance with Duke Institutional Animal Care and Use Committee and the EC Directive on the protection of animals used for scientific purposes. Animals were housed on a 12 hour light/dark cycle and provided food and water *ad libitum*.

### Breeding and Maintenance of Gz-knockout mouse line

Two 7-week-old male Gz+/-Balb/cAnNcrIBR (Balb/c) mice were obtained in January 2005 from Ian Hendry at Australian National University. These mice had been back-crossed by the Hendry group into the Balb/c background for over 10 generations and were mated with 6-week-old female Balb/c mice from Charles River Labs to generate founders for the Duke University colony. The colony was maintained using Het x Het breeding triads of one male and two females. Breeding triads were replaced when litter size declined (at approximately 9-12 months of age) a total of seven times throughout the course of these studies over five years. New breeding triads were chosen from offspring of different litters to limit any genetic drift. Approximately three experiments were conducted per generation. Whenever possible, wild-type littermate controls were used for the pilocarpine seizure experiments; otherwise, age and weight-matched animals were chosen such that 54% of the wild-type animals used in these studies were littermate controls. Notably, when housed with wild-type mice, Gz−/−Balb/c mice routinely gain excessive weight; therefore, experimental mice were segregated by genotype at 3 weeks of age (prior to experiments).

### Pilocarpinecarpine-induced seizure studies

Sixteen week-old male mice in an inbred BALB/c background were used to characterize seizure behavior. All mice were pre-treated with a 2 mg/kg dose of scopolamine methyl-bromide and terbutaline hemisulfate (Sigma) by intraperitoneal (IP) injection to prevent peripheral effects of the pilocarpine. After 30 min, mice were treated with pilocarpine hydrochloride (Sigma) by IP injection. Mice were observed and their behavior was recorded in 5 min windows for 4 h. Behavior was classified as follows: stage 0, no behavioral sign of seizure; stage 1, facial stereotypes such as lip smacking or foaming at the mouth, as well as loss of conscious response to sounds and smells; stage 2, compulsive head nodding; stage 3, forelimb clonus (rapid shuffling of paws); stage 4, seizure including rearing onto hind limbs; stage 5, seizures involving barrel rolling; FS, fatal SE, seizure terminating in death. Status epilepticus was defined as a stage 2 or higher lasting for at least 30 minutes or longer.

### Electrode Design and Surgery

Recording electrodes composed of Tungsten wires (50µm) arranged into bundles were surgically implanted into the brain to gather LFP recordings as described by Dzirasa, et al. ^54^. The following brain regions were implanted with the corresponding coordinates, anterior/posterior (A/P) and medial/lateral (M/L), measured in millimeters (mm) from Bregma, and dorsal/ventral (D/V) measured from brain: amygdala (−1.3 A/P, 3.0 M/L, −4.6D/V), dorsal hippocampus (−2.2 A/P, 1.5-2.25 M/L, −2.0D/V), ventral hippocampus (3.0 A/P, 3.0 M/L, − 2.5D/V), thalamus (−0.6 A/P, 0.6 M/L, −3.5D/V), motor (2.1 A/P, 2.0 M/L, −1.0D/V) and sensory cortices (.02 A/P, 2.7 M/L, −1.6D/V). Briefly, mice were treated with anesthetics, ketamine (100mg/kg) and xylazine (10mg/kg), and placed on a stereotaxic device where the scalp was cut at the midline to expose the skull. Three ground screws were embedded in the front, back, and to the right of the skull.

### Electrophysiological Recordings

For the electrophysiology experiments, four animals per group (Gz-null and WT cohorts) were implanted. After surgery, all mice were habituated to being handled and moving while being plugged in to the electrophysiological recordings apparatus. The recording apparatus was constructed such that the mice were able to move freely while plugged in. During electrophysiological recordings, mice were connected to a recording headstage and LFP data was collected at 1000Hz using Blackrock Microsystem CerePlex™ Direct acquisition equipment and software. After the habituation period, mice were treated with pilocarpine, in the same manner as for the behavioral experiments, and electrophysiological recordings were collected throughout the duration of status epilepticus.

### Electrographic Seizure Identification and Data Analysis

MATLAB was used to create visual graphics of LFP recordings as voltage (mV) against time (milliseconds). An electrographic seizure was defined as a consistent, irregular set of waveforms that lasted for at least 10 seconds ^55^ (Figure 2). Also, the time of the first seizure lasting 10 seconds was identified as seizure onset as has been done in various studies ^42,55,56^.

## Acknowledgements

We thank Debra J. Evanson and Missy Infante for technical assistance. We thank the Laboratory of James McNamara, A. Soren Leonard, Sarah Jacobs, Michelle Kimple, Ian Cushman, James Crowley and J.H. Pate Skene for helpful discussion and insight.

## Abbreviations used

BDNF: brain-derived neurotrophic factor
CNS: central nervous system
NE: norepinephrin
DA: dopamine
5-HT: serotonin

## References

1. Fisher, R. S. et al. Epileptic seizures and epilepsy: definitions proposed by the International League Against Epilepsy (ILAE) and the International Bureau for Epilepsy (IBE). Epilepsia 46, 470–472 (2005).

2. Annegers, J. F. & Coan, S. P. SUDEP: overview of definitions and review of incidence data. Seizure 8, 347–352 (1999).

3. Tsai, S.-J. Is autism caused by early hyperactivity of brain-derived neurotrophic factor?. Medical hypotheses 65, 79–82 (2005).

4. Leonardi, M. & Ustun, T. B. The global burden of epilepsy. Epilepsia 43 Suppl 6, 21–25 (2002).

5. Binder, D. K. in Recent Advances in Epilepsy Research 34–56 (Springer, 2004).

6. McNamara, J. O., Huang, Y. Z. & Leonard, A. S. Molecular signaling mechanisms underlying epileptogenesis. Sci STKE 2006, re12 (2006).

7. Ernfors, P., Bengzon, J., Kokaia, Z., Persson, H. & Lindvall, O. Increased levels of messenger RNAs for neurotrophic factors in the brain during kindling epileptogenesis. Neuron 7, 165–176, doi:0896-6273(91)90084-D [pii] (1991).

8. Gall, C. M. & Isackson, P. J. Limbic seizures increase neuronal production of messenger RNA for nerve growth factor. Science 245, 758–761 (1989).

9. Isackson, P. J., Huntsman, M. M., Murray, K. D. & Gall, C. M. BDNF mRNA expression is increased in adult rat forebrain after limbic seizures: temporal patterns of induction distinct from NGF. Neuron 6, 937–948, doi:0896-6273(91)90234-Q [pii] (1991).

10. Nibuya, M., Morinobu, S. & Duman, R. S. Regulation of BDNF and trkB mRNA in rat brain by chronic electroconvulsive seizure and antidepressant drug treatments. The Journal of neuroscience: the official journal of the Society for Neuroscience 15, 7539–7547 (1995).

11. Takahashi, M. et al. Patients with temporal lobe epilepsy show an increase in brain-derived neurotrophic factor protein and its correlation with neuropeptide Y. Brain research 818, 579–582, doi:S0006-8993(98)01355-9 [pii] (1999).

12. Kokaia, M. et al. Suppressed epileptogenesis in BDNF mutant mice. Exp Neurol 133, 215–224, doi:S0014-4886(85)71024-2 [pii] 10.1006/exnr.1995.1024 (1995).

13. He, X. P. et al. Conditional deletion of TrkB but not BDNF prevents epileptogenesis in the kindling model. Neuron 43, 31–42 (2004).

14. Croll, S. D. et al. Brain-derived neurotrophic factor transgenic mice exhibit passive avoidance deficits, increased seizure severity and in vitro hyperexcitability in the hippocampus and entorhinal cortex. Neuroscience 93, 1491–1506, doi:S0306-4522(99)00296-1 [pii] (1999).

15. Scharfman, H. E., Goodman, J. H., Sollas, A. L. & Croll, S. D. Spontaneous limbic seizures after intrahippocampal infusion of brain-derived neurotrophic factor. Exp Neurol 174, 201–214, doi:10.1006/exnr.2002.7869 S0014488602978696 [pii] (2002).

16. Hultman, R., Kumari, U., Michel, N. & Casey, P. J. Gαz regulates BDNF-induction of axon growth in cortical neurons. Molecular and Cellular Neuroscience 58, 53–61 (2014).

17. Meng, J., Glick, J. L., Polakis, P. & Casey, P. J. Functional interaction between Galpha(z) and Rap1GAP suggests a novel form of cellular cross-talk. J Biol Chem 274, 36663–36669 (1999).

18. Embry, A. C., Glick, J. L., Linder, M. E. & Casey, P. J. Reciprocal signaling between the transcriptional co-factor Eya2 and specific members of the Galphai family. Mol Pharmacol 66, 1325–1331, doi:10.1124/mol.104.004093 (2004).

19. Glick, J. L., Meigs, T. E., Miron, A. & Casey, P. J. RGSZ1, a Gz-selective regulator of G protein signaling whose action is sensitive to the phosphorylation state of Gzalpha. J Biol Chem 273, 26008–26013 (1998).

20. Casey, P. J., Fong, H. K., Simon, M. I. & Gilman, A. G. Gz, a guanine nucleotide-binding protein with unique biochemical properties. J Biol Chem 265, 2383–2390 (1990).

21. Gagnon, A. W. et al. Identification of Gz alpha as a pertussis toxin-insensitive G protein in human platelets and megakaryocytes. Blood 78, 1247–1253 (1991).

22. Kimple, M. E., Hultman, R., Casey, P.J. in Handbook of Cell Signaling Vol. 2 1649–1653 (Academic Press, 2010).

23. Ho, M. K. & Wong, Y. H. Structure and function of the pertussis-toxin-insensitive Gz protein. Biol Signals Recept 7, 80–89 (1998).

24. Ho, M. K. & Wong, Y. H. G(z) signaling: emerging divergence from G(i) signaling. Oncogene 20, 1615–1625, doi:10.1038/sj.onc.1204190 (2001).

25. Oleskevich, S., Leck, K.-J., Matthaei, K. & Hendry, I. A. Enhanced serotonin response in the hippocampus of Gαz protein knock-out mice. Neuroreport 16, 921–925 (2005).

26. van den Buuse, M., Martin, S., Holgate, J., Matthaei, K. & Hendry, I. Mice deficient in the alpha subunit of G(z) show changes in pre-pulse inhibition, anxiety and responses to 5-HT(1A) receptor stimulation, which are strongly dependent on the genetic background. Psychopharmacology (Berl) 195, 273–283, doi:10.1007/s00213-007-0888-7 (2007).

27. Leck, K. J. et al. Gz proteins are functionally coupled to dopamine D2-like receptors in vivo. Neuropharmacology 51, 597–605, doi:10.1016/j.neuropharm.2006.05.002 (2006).

28. Kelleher, K. L., Matthaei, K. I. & Hendry, I. A. Targeted disruption of the mouse Gz-alpha gene: a role for Gz in platelet function?. Thromb Haemost 85, 529–532 (2001).

29. Yang, J. et al. Loss of signaling through the G protein, Gz, results in abnormal platelet activation and altered responses to psychoactive drugs. Proceedings of the National Academy of Sciences of the United States of America 97, 9984–9989, doi:10.1073/pnas.180194597 (2000).

30. Garzon, J., Martinez-Pena, Y. & Sanchez-Blazquez, P. Gx/z is regulated by mu but not delta opioid receptors in the stimulation of the low Km GTPase activity in mouse periaqueductal grey matter. The European journal of neuroscience 9, 1194–1200 (1997).

31. Kimple, M. E. et al. Deletion of GalphaZ protein protects against diet-induced glucose intolerance via expansion of beta-cell mass. J Biol Chem 287, 20344–20355, doi:10.1074/jbc.M112.359745 (2012).

32. van den Buuse, M., van Driel, I. R., Samuelson, L. C., Pijnappel, M. & Martin, S. Reduced effects of amphetamine on prepulse inhibition of startle in gastrin-deficient mice. Neurosci Lett 373, 237–242, doi:10.1016/j.neulet.2004.10.013 (2005).

33. van den Buuse, M. et al. Enhanced effect of dopaminergic stimulation on prepulse inhibition in mice deficient in the alpha subunit of G(z). Psychopharmacology (Berl) 183, 358–367, doi:10.1007/s00213-005-0181-6 (2005).

34. Cavalheiro, E. A., Fernandes, M. J., Turski, L. & Naffah-Mazzacoratti, M. G. Spontaneous recurrent seizures in rats: amino acid and monoamine determination in the hippocampus. Epilepsia 35, 1–11 (1994).

35. Tecott, L. H. et al. Eating disorder and epilepsy in mice lacking 5-HT2c serotonin receptors. Nature 374, 542–546, doi:10.1038/374542a0 (1995).

36. Vriend, J., Alexiuk, N. A., Green-Johnson, J. & Ryan, E. Determination of amino acids and monoamine neurotransmitters in caudate nucleus of seizure-resistant and seizure-prone BALB/c mice. J Neurochem 60, 1300–1307 (1993).

37. Svob Strac, D. et al. Monoaminergic Mechanisms in Epilepsy May Offer Innovative Therapeutic Opportunity for Monoaminergic Multi-Target Drugs. Front Neurosci 10, 492, doi:10.3389/fnins.2016.00492 (2016).

38. Kandratavicius, L. et al. Animal models of epilepsy: use and limitations. Neuropsychiatr Dis Treat 10, 1693–1705, doi:10.2147/NDT.S50371 (2014).

39. Curia, G., Longo, D., Biagini, G., Jones, R. S. & Avoli, M. The pilocarpine model of temporal lobe epilepsy. Journal of neuroscience methods 172, 143–157, doi:10.1016/j.jneumeth.2008.04.019 (2008).

40. Pitkanen, A. Models of Seizures and Epilepsy. (Academic Press, 2005).

41. Maganti, R. K. & Rutecki, P. EEG and epilepsy monitoring. Continuum (Minneap Minn) 19, 598–622, doi:10.1212/01.CON.0000431378.51935.d8 (2013).

42. Toyoda, I., Bower, M. R., Leyva, F. & Buckmaster, P. S. Early activation of ventral hippocampus and subiculum during spontaneous seizures in a rat model of temporal lobe epilepsy. The Journal of neuroscience: the official journal of the Society for Neuroscience 33, 11100–11115, doi:10.1523/JNEUROSCI.0472-13.2013 (2013).

43. Rosenbloom, K. R. et al. The UCSC Genome Browser database: 2015 update. Nucleic Acids Res 43, D670–681, doi:10.1093/nar/gku1177 (2015).

44. Lindgren, V. et al. Behavioral abnormalities are common and severe in patients with distal 22q11.2 microdeletions and microduplications. Mol Genet Genomic Med 3, 346–353, doi:10.1002/mgg3.146 (2015).

45. Gaitatzis, A., Trimble, M. & Sander, J. W. The psychiatric comorbidity of epilepsy. Acta Neurologica Scandinavica 110, 207–220 (2004).

46. Rocha, L. et al. Do certain signal transduction mechanisms explain the comorbidity of epilepsy and mood disorders?. Epilepsy Behav 38, 25–31, doi:10.1016/j.yebeh.2014.01.001 (2014).

47. Binder, D. K. The role of BDNF in epilepsy and other diseases of the mature nervous system. Adv Exp Med Biol 548, 34–56 (2004).

48. Pericic, D., Lazic, J., Jazvinscak Jembrek, M. & Svob Strac, D. Stimulation of 5-HT 1A receptors increases the seizure threshold for picrotoxin in mice. Eur J Pharmacol 527, 105–110, doi:10.1016/j.ejphar.2005.10.021 (2005).

49. Toth, M. 5-HT1A receptor knockout mouse as a genetic model of anxiety. Eur J Pharmacol 463, 177–184 (2003).

50. Ramboz, S. et al. Serotonin receptor 1A knockout: an animal model of anxiety-related disorder. Proceedings of the National Academy of Sciences of the United States of America 95, 14476–14481 (1998).

51. Clinckers, R., Smolders, I., Meurs, A., Ebinger, G. & Michotte, Y. Anticonvulsant action of GBR-12909 and citalopram against acute experimentally induced limbic seizures. Neuropharmacology 47, 1053–1061, doi:10.1016/j.neuropharm.2004.07.032 (2004).

52. Parmentier, M. L. et al. The G protein-coupling profile of metabotropic glutamate receptors, as determined with exogenous G proteins, is independent of their ligand recognition domain. Mol Pharmacol 53, 778–786 (1998).

53. Kammermeier, P. J., Davis, M. I. & Ikeda, S. R. Specificity of metabotropic glutamate receptor 2 coupling to G proteins. Mol Pharmacol 63, 183–191 (2003).

54. Dzirasa, K., Fuentes, R., Kumar, S., Potes, J. M. & Nicolelis, M. A. Chronic in vivo multi-circuit neurophysiological recordings in mice. Journal of neuroscience methods 195, 36–46 (2011).

55. Bragin, A., Azizyan, A., Almajano, J., Wilson, C. L. & Engel, J., Jr. Analysis of chronic seizure onsets after intrahippocampal kainic acid injection in freely moving rats. Epilepsia 46, 1592–1598, doi:10.1111/j.1528-1167.2005.00268.x (2005).

56. Bower, M. R. & Buckmaster, P. S. Changes in granule cell firing rates precede locally recorded spontaneous seizures by minutes in an animal model of temporal lobe epilepsy. J Neurophysiol 99, 2431–2442, doi:10.1152/jn.01369.2007 (2008).

